# Comprehensive analysis of human microRNA-mRNA interactome

**DOI:** 10.1101/675694

**Authors:** Olga Plotnikova, Ancha Baranova, Mikhail Skoblov

**Affiliations:** Laboratory of functional genome analysis, Moscow Institute of Physics and Technology, Moscow, Russia; Laboratory of functional genomics, Research Centre for Medical Genetics, Moscow, Russia; School of Systems Biology, George Mason University, Fairfax, VA, USA; School of Biomedicine, Far Eastern Federal University, Vladivostok, Russia

**Keywords:** microRNA, regulation of gene expression, microRNA-mRNA interactions, microRNA binding sites, miRNA-target RNA duplexes, web tool for searching microRNA-binding regions

## Abstract

MicroRNAs play a key role in the regulation of gene expression. A majority of microRNA-mRNA interactions remain unidentified. Despite extensive research, our ability to predict human microRNA-mRNA interactions using computational algorithms remains limited by a complexity of the models for non-canonical interactions, and an abundance of false positive results.

Here we present the landscape of microRNA-mRNA human interactions, which we derived from comprehensive analysis of datasets describing direct microRNA-mRNA interactions experimentally defined in HEK293 and Huh7.5 cell lines, along with other available microRNA and mRNA expression data. We have also established a collection of reliable microRNA binding regions that we systematically extracted in course of analysis of 79 CLIP datasets, which is available at http://score.generesearch.ru/services/mirna/.

While only 1-2% of human genes interact with microRNAs, some RNAs display a substantial sponge effect, which is specific to the cell line of study. Some microRNAs are expressed at a very high level, while interacting with only a few mRNAs, thus, indeed, serving as specific gene expression regulators. Other miRNAs might be expressed at relatively low levels, and interact with many mRNAs. Some of the microRNAs might switch between these two classes, depending on cellular context. Results of our study provide an initial resolution into the complex patterns of human microRNA-mRNA interactions.

## 1 Introduction

MicroRNAs are small noncoding RNAs that associate with Argonaute (*AGO*) protein to form a silencing complex, which then regulates a gene expression (Jonas and Izaurralde, 2015). MicroRNAs accomplish essential post-transcriptional regulatory step of gene expression regulation through ether the degradation of a transcript or the inhibition of translation, and are involved in key cellular processes, such as apoptosis, proliferation or differentiation (He and Hannon, 2004). Hence, the dysregulation of microRNAs could result in the development of a disease or in a malignant transformation (Weiss and Ito, 2017). According to some estimates, nearly all mature sequences of coding transcripts contain potential sites for microRNA regulation (Bartel, 2004; Friedman et al., 2009).

Human genome encodes approximately 2600 mature microRNAs (miRBase v.22) and, according to GENCODE data (v.29), more than 200 thousands of transcripts, including isoforms with slight variations. A particular microRNA may target many different mRNAs (Selbach et al., 2008); a particular messenger RNA may bind to a variety of microRNAs, either simultaneously or in context-dependent fashion (Uhlmann et al., 2012). Notably, within some messenger RNAs, the target regions for particular microRNAs cluster together, resulting in the cooperative repression effect (Grimson et al., 2007; Sætrom et al., 2007). The mapping of microRNA-mRNA interactions is far from being complete due to the recognized challenges of computational prediction of mRNA-microRNA interactions.

In our previous study, we showed that the outputs generated by commonly used microRNA-mRNA interactions predicting software differ from each other substantially, while failing correctly pinpoint microRNA-binding regions identified in wet lab experiments (Plotnikova and Skoblov, 2018). Nowadays, many tools for the prediction microRNA-mRNA interactions are in development, all with different underlying algorithms (Riffo-Campos et al., 2016; Gumienny and Zavolan, 2015; Lu and Leslie, 2016; Agarwal et al., 2015;). Among most advanced algorithms we should highlight the ones taking into account expression levels of both the microRNAs and their targets. Notably, the changes in expression of microRNA may also affect expression levels of other, non-target mRNAs, for example, due miRNA targeting of their upstream regulators. Consequently, newer, more comprehensive approaches, like miRImpact (Artcibasova et al., 2016), PanMiRa (Li and Zhang, 2014), and ProMISe (Li et al., 2014), aim at explaining complex phenotypes by performing analysis of each microRNAs along with its direct and indirect targets.

The experimental identification of direct microRNA targets remains a crucial step in attaining good prediction results. There are two main groups of the experimental approaches for a direct identification of microRNA-mRNA interactions. The first approach relies on a construction of reporter gene assays and one-by-one evaluation of possible interactions between the microRNA and its cognate mRNA region of interest through measuring the activity of the reporter (Steinkraus et al., 2016). Another group of techniques comprises involves a coupling of a cross-linking with immunoprecipitation (CLIP); this group represented by variety of the protocols including PAR-CLIP, iCLIP, HITS-CLIP, and others (Steinkraus et al., 2016; Licatalosi et al., 2008). CLIP group of methods identifies the microRNA binding regions in target mRNAs only, while information about pairing of a particular microRNA with a particular mRNA region remains obscure.

There are two modifications of AGO-CLIP based technology developed specifically for identifying microRNAs ligated to their endogenous mRNA targets as part of chimeric molecules. To date, evaluations of microRNA-mRNA interactomes by these two technologies utilized only two human cell lines. Helwak and colleagues applied so-called cross-linking ligation and sequencing of hybrids, or CLASH, to HEK293 cell line, retrieving more than 18,000 high-confidence microRNA-mRNA interactions (Helwak et al., 2013). Later, Moore and colleagues used another variety of AGO-CLIP termed CLEAR (covalent ligation of endogenous Argonaute-bound RNAs)-CLIP for the study of microRNA-interactome in Huh7.5 cell (Moore et al., 2015). CLASH and CLEAR-CLIP techniques closely resemble each other, with the only difference that CLASH protocol employs HEK293 cell line over-expressed AGO1, while CLEAR-CLIP targets endogenous AGO allowing experimenting with any cell line. Thus, CLEAR-CLIP does not require full denaturation of AGO and involves a single purification step. It is of note that both publications cited above concentrated on the development of the experimental protocol and subsequent evaluation of the technical aspects of analytic procedure, rather than on extracting biological insights from the data collected.

We aggregated various experimental data on human miRNA-mRNA interactions, and analyzed them. First, we investigate how expression levels of microRNAs and their cognate mRNAs correlate, and if the behavior of miRNA-mRNA pairs depends on a cell line context. In order to do this, we analyzed together (i) sequences and abundance of microRNA and their target mRNAs in CLASH dataset for HEK293 cell line and in CLEA-CLIP dataset for Huh7.5 cell line and (ii) expression level of microRNAs and RNAs in HEK293 and in Huh7.5 cell lines. Second, we attempted an identification of a credible, experimentally confirmed microRNA binding regions in CLASH/ CLEAR-CLIP datasets and in 79 additional CLIP datasets.

## 2 Materials and Methods

### 2.1 microRNA-mRNA interactions

microRNA-mRNA interactome data were extracted from published CLASH (Helwak et al., 2013) and CLEAR-CLIP (Moore et al., 2015) studies. Using Ensemble API, the coordinates of microRNA – mRNA interacting regions were transformed into genome coordinates. In total, we revealed 18,478 microRNA-mRNA interactions in 22,030 genome regions. For a total of 36 interactions, the transforming of their coordinates failed. We used LiftOver to transform CLEAR-CLIP interactome data from hg18 genome version into hg19. wAnnovar (Wang et al., 2010; Yang and Wang, 2015) was used to annotate genomic regions (CDS, 3’UTR, 5’UTR, intronic, intergenic, etc). To estimate the expected overlap between CLASH and CLEAR-CLIP like datasets we used a custom python script.

### 2.2 mRNA expression

Publicly available RNA-seq datasets GSE68611 (Murakawa et al., 2015) and GSE64677 (Luna et al., 2015) were used for extracting and examining gene sets expressed in HEK293 and Huh7.5 cell lines. Each of these datasets includes two biological replicates. Initial quality control of sequencing outputs was performed using FastQC. Next, we used kallisto (Bray et al., 2016) to map raw reads to the human reference transcript sequences (GENCODE, 28 version).

First, in each experiment, we calculated the gene expression levels as the sum of expression levels for individual gene transcripts. Second, we took the mean value for each gene between two processed datasets in each of the two cell lines. Finally, we kept only genes that had expression more or equal to 1 tpm as total value and that had expression level of at the level at least 1 tpm in one of the two experiments.

To reveal an amount of interactions with microRNAs for genes, we used CLASH and CLEAR-CLIP datasets for HEK293 and Huh7.5 cell lines, respectively.

Gene functions were interpreted using PANTHER toolkit Version 12.0 (http://www.pantherdb.org/tools). We used InteractiVenn tool (Heberle et al., 2015) to create Venn diagrams in our analysis.

### 2.3 microRNA expression

We downloaded microRNA expression data from the GEO database: two experimental replicates for HEK293 cell line (GSE75136 (Wissink et al., 2016)) and three experimental replicates for Huh7.5 cell line (GSE74014 (Bandiera et al., 2016)). The correlations of experimental results obtained in two cell lines were calculated using the Spearman’s procedure. We used the R package “DeSeq2” to normalize microRNA expression. MicroRNA was considered as expressed if it had expression more than 3 counts.

CLASH and CLEAR-CLIP datasets were used to calculate the amount of interactions for each microRNAs. The correlation of the amounts of interactions formed by microRNAs and their expression levels were estimated using the Spearman correlation coefficient.

In order to calculate a conservative phyloP score for all microRNAs we downloaded the coordinates of the mature microRNAs from miRBase (Kozomara and Griffiths-Jones, 2013) (release 22, coordinates corresponded to the GRCh38 human reference genome). Next, we used UCSC table browser (Karolchik et al., 2004) to obtain phyloP conservative values across 20 vertebrates for all mature microRNAs. For each group of microRNAs, the mean value between the phyloP scores was calculated.

### 2.4 CLIP-data

We collected 79 CLIP datasets (Supplementary Table 3) from the POSTAR database (Hu et al., 2016) that were initially preprocessed by unified procedures: PAR-CLIP datasets (N = 18) by PARalyzer (Corcoran et al., 2011) method and HITS-CLIP datasets (N = 61) by CIMS (Moore et al., 2014) method. We used python to analyze all microRNA binding regions from CLIP datasets together with microRNA-mRNA interactions from CLASH and CLEAR-CLIP. In total, all regions were merged in six million nucleotides and each position was characterized by the following parameters: list of supported experiments (GEO GSM ID), their corresponding cell lines and list of interacted microRNAs (if accessible). We used wAnnovar to annotate genes and their parts (CDS, 3’UTR, 5’UTR, intronic, etc).

### 2.5 microRNA binding regions

Our analysis of CLIPs, CLASH, and CLEAR-CLIP revealed 156 thousand regions. We used a custom python script to select experimentally confirmed microRNA binding regions (Exp-MiBR). Exp-MiBR was defined as a region that had a subsequence of length L=10, whereas each nucleotide (position) in this subsequence had been supported by at least n=2 different datasets or chimeras. We estimated the amount of Exp-MiBRs for all combination of length and amount of supported datasets/chimeras in ranges: L=1-25 and n=1-10 (Supplementary Table 5).

### 2.6 Exp-MiBRs application

We characterized each Exp-MiBR (total amount = 46805) by the following parameters: gene information; amount and list of supported experiments (GEO GSM ID) and their corresponding cell lines; list of interacted microRNAs (if accessible).

Besides that all the Exp-MiBRs with the corresponded information are available as Supplementary Table 4, we also provide an open-access web tool via http://score.generesearch.ru/services/mirna. As input, the tool requires any VCF file (v4.0 or 4.1), no more than 20MB or a single (point) genome coordinate. The file or coordinate could be recorded in human genome assembly version 38 or 19.

### 2.7 Web tool for searching Exp-MiBRs

All microRNA binding regions identified as experimentally confirmed (Exp-MiBR) and reported in this paper (Supplementary Table 4) may be searched by a web tool available online: http://score.generesearch.ru/services/mirna/.

## 3 Results

### 3.1 Comparison of high-throughput microRNA-mRNA interactions from CLASH and CLEAR-CLIP datasets

First, we compared the sets of microRNA-mRNA interactions retrieved in HEK293 and in Huh7.5 by CLASH (Helwak et al., 2013) and CLEAR-CLIP (Moore et al., 2015) protocol, respectively. Although CLASH and CLEAR-CLIP techniques are somewhat similar, CLEAR-CLIP study (N=32,170) revealed almost two times more interactions than CLASH study (N=18,478). One of the reasons for this may be due to the differences in the data processing procedures. While CLASH sequences were aligned to the mature transcriptome, CLEAR-CLIP data have been mapped to human genome. Because of that, CLEAR-CLIP technique was capable to highlight additional interaction sites located in the introns and the intergenic regions (∼70% of all interactions).

To enable the comparison, we focused our analysis on miRNA binding regions residing within the mature transcriptome. Because of that, CLEAR-CLIP dataset was limited to about one-third of its entries (n=10,032). Further analysis estimated that approximately 2-3% of the total length of all expressed protein-coding transcripts serve as a target for one or another microRNAs in either CLASH or CLEAR-CLIP datasets. In addition, in both datasets, the microRNA binding regions had similar distribution by mRNA regions (3’UTR, CDS, 5’UTR), and to the distribution of the mRNA parts present in GENCODE (Figure 1A). Thus, the datasets generated by CLASH and CLEAR-CLIP techniques are comparable.

**Figure 1.**
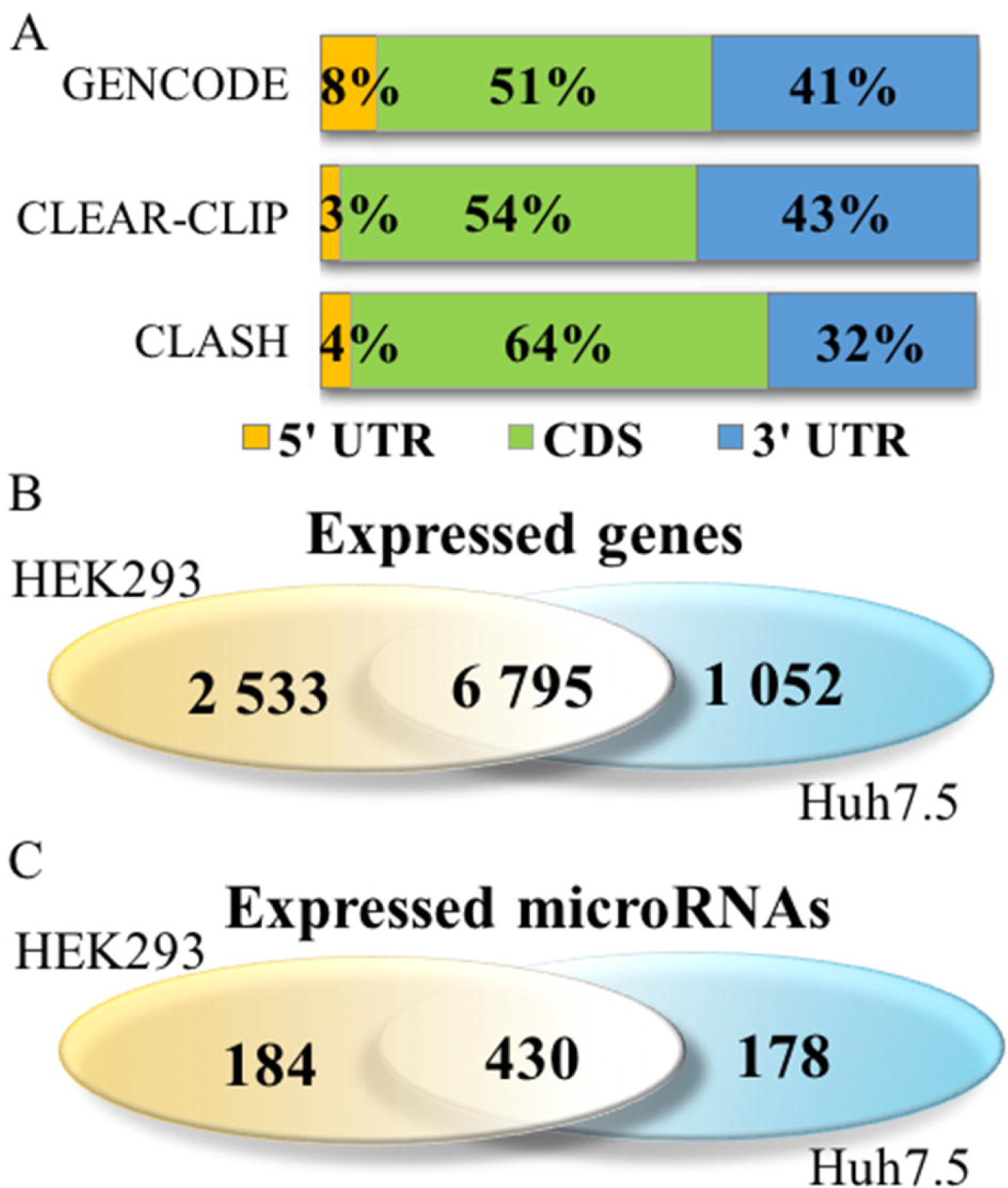
A comparison of CLASH and CLEAR-CLIP datasets. **(A)** Distribution of the summarized lengths of 3’UTR, CDS or 5’UTR mRNA regions in CLEAR-CLIP, CLASH and GENCODE, respectively. **(B)** Venn diagram of HEK293- and Huh7.5-expressed genes as covered by CLASH and CLEAR-CLIP interactomes, respectively. **(C)** Venn diagram of HEK293- and Huh7.5-expressed miRNAs represented in CLASH and CLEAR-CLIP interactomes, respectively.

Comparison of these two studies revealed approximately one thousand common binding regions found both in a set of eighteen thousand interactions from CLASH and in a set of ten thousand interactions from CLEAR-CLIP. To evaluate if this overlap reflect biological phenomenon rather than statistical fluke, we performed computational simulation of CLASH and CLEAR-CLIP interactions in transcripts expressed in HEK293 (N = 7,299) and Huh7.5 (N = 4,977), respectively. For these cell lines, a common set of expressed mRNAs (n = 3,044) was reduced to a set of randomly selected nucleotide fragments with the size distribution matching that for nucleotide fragments of CLASH and CLEAR-CLIP, then we analyzed these sets of sequences for overlap. After five independent runs with randomly selected fragments of matching size distribution, we detected, on average, 7.4 +/-1.3 interactions with an average length of overlapped segments at 14 nt +/-6.7 nt. Among these interactions, only a fraction had the length of overlap of more than 20 nt (5.0 +/-2.5). In the experimentally obtained CLASH and CLEAR-CLIP datasets, we detected 1,153 common miRNA-mRNA interactions, built upon combinations of 933 fragments interacting in CLASH and 944 fragments interacting in CLEAR-CLIP. Average length of experimentally obtained interaction was at 37.2 nt +/-19.4 nt. Eight hundred and sixty seven interactions which were common for both datasets had the length of overlap of more than 20 nt, with an average length of 45.8 nt +/-13.9 nt. Therefore, the characteristics of experimentally detected patterns of miRNA-mRNA interactions differ from that of interactions generated by simulation of random events (P < 0.0001).

To investigate whether the low degree of the overlap between miRNA-mRNA interactions registered in CLASH and CLEAR-CLIP datasets could be due to low degree of the overlap between HEK293 and Huh7.5 transcriptomes, expression data collected from these two cell lines were downloaded from GEO repository and analyzed. While about half of expressed microRNAs were common for both these cell lines (Figure 1C), overall difference in expression patterns of HEK293 and Huh7.5 cells (Figure 1B) was clearly evident. To find out if cell-specific differences in microRNA-mRNA interactomes are due to cell-specific environment, the relationships between the levels of expression for individual miRNAs and their targets as well as the patterns of interactions for each mRNAs and miRNAs in the both cell lines were investigated in details.

### 3.2 Expression analysis of microRNA-mRNA interactome

#### 3.2.1 mRNA expression analysis

To investigate the degree to which cell-specific levels of transcripts depend on respective microRNAs, we compared expression levels of each gene in HEK293 and Huh7.5 cell lines, then cross-compared them to sets of experimentally detected microRNA interactions. HEK293 and Huh7.5 cell lines express a total of 15,8k and 14,5k genes, respectively. In each of these two cell lines, approximately 6.9k genes interacted with one or more microRNAs (Supplementary Fig. 1). Our analysis pinpointed 1-2% of mRNAs with confirmed interactions and no expression detected in the corresponding cell line. It is possible that these mRNAs have been detected as chimeric reads resulting from their protection by AGO protein from Ribonucleases. Below, we will describe a few microRNAs that were detected only as a part of chimeras.

In each of these cell lines, a majority of expressed mRNAs (57-59%) did not interact with any microRNA (Figure 2AB). In CLASH and CLEAR-CLIP datasets, there were 215 and 333 high-interacting mRNAs, respectively, with nine or more miRNA interactions for each.

**Figure 2.**
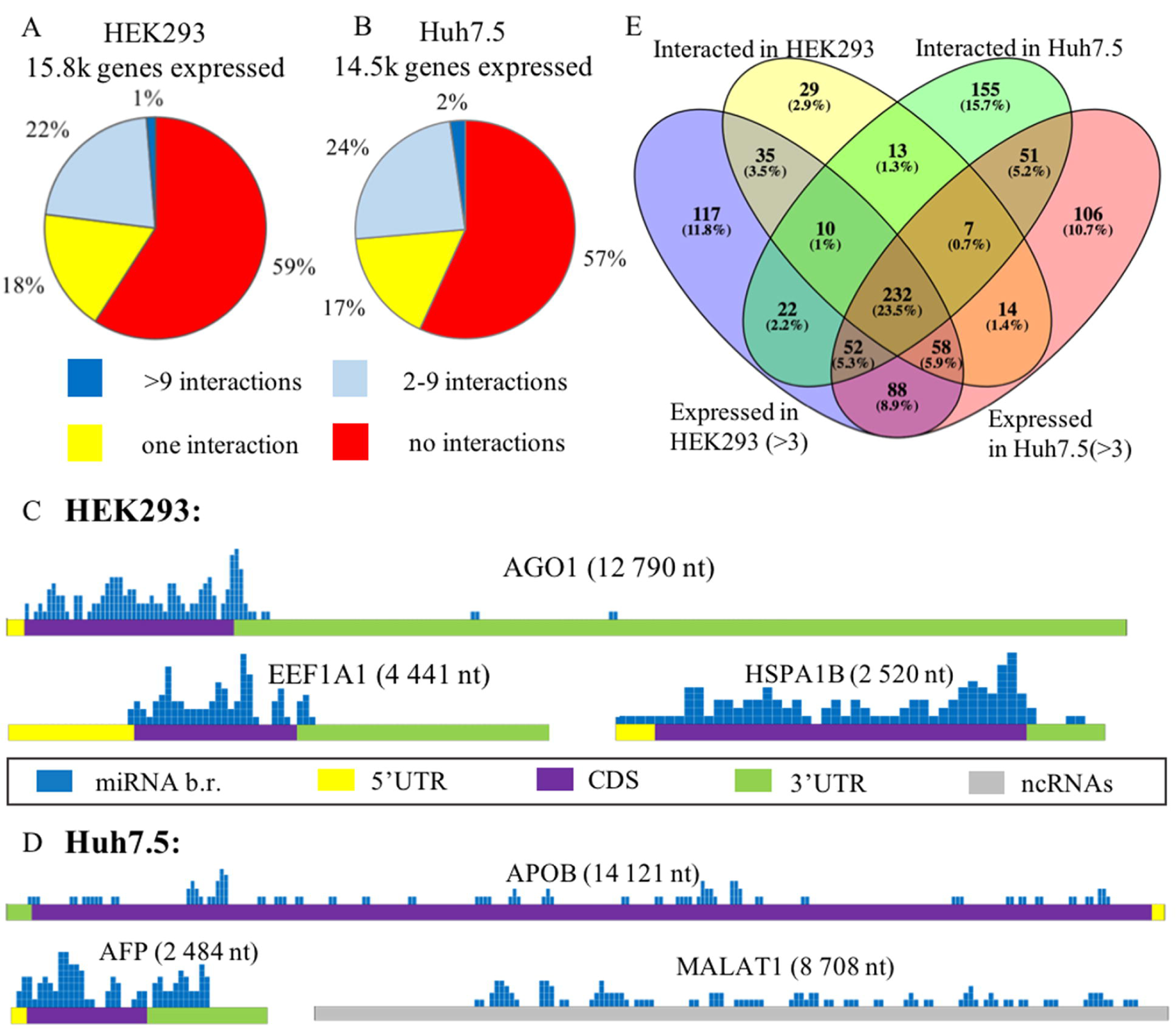
microRNA and mRNA expression analysis in HEK293 and Huh7.5 cell lines. **(A) and (B)**: Analysis of expressed genes according to amounts of their interactions with microRNAs in HEK293 **(A)** and Huh7.5 **(B)** cell lines; **(C)** and **(D)**: Locations of experimentally confirmed microRNA binding regions (Exp-MiBRs) in sponge-like RNAs expressed in HEK293/CLASH **(C)** and Huh7.5/ CLEAR-CLIP **(D)** datasets. After segmenting each of the presented RNAs into 50nt-pieces, the segments coinciding with Exp-MiBR were marked blue on the mRNA map. For each sponge RNA, name and length are above the gene schematics. Colored parts of RNAs are as follows: 5’UTR –yellow, coding region – violet, 3’UTR – green, noncoding region – grey. **(E)** The overlaps between expressed and interacting microRNAs in HEK293 and Huh7.5 cell lines.

Cell line-specific pie charts built for the miRNA-mRNA interactions per mRNAs were similar. Nevertheless, comparison of the most regulated sets of genes with 9 or more interactions each revealed that these sets were cell-line-specific, with only 18 genes in common. These common eighteen genes formed in average of 15.7+/-3.2 and 14.1+/-2.4 interactions with microRNAs in the HEK293 and Huh7.5 cell lines, respectively. Surprisingly, cell line-specific sets of microRNA regulators for each of these genes were completely different. By PANTHER analysis of the common set of genes, we detected enrichment in only one Gene Ontology (GO) category – a molecular function of RNA binding (Supplementary Table 1).

Further, we identified a set of mRNAs capable of interaction with many different types of microRNA molecules, with no preference to a particular miRNA. Such behavior of ambigious interaction with many microRNAs is similar to “sponge” performance of circular RNAs and lncRNAs. Among “sponge-like” mRNAs with 50 or more interactions detected for each were *AGO1, EEF1A1* and *HSPA1B* in HEK293/CLASH. Peculiarly, in Huh7.5/ CLEAR-CLIP, same property attributed to different set of mRNA, namely, *APOB, AFP, MALAT1* and *XIST*. In mRNAs with sponge-like property, microRNA interaction sites were located predominantly in the protein coding part (Figure 2C and 2D).

Remarkably, in HEK293 cells, the most interacting mRNA was the one for AGO1 protein, which had been overexpressed on purpose, as part of CLASH protocol. In this cell line, AGO1-encoding mRNA has 88 interactions with a total of 50 different microRNAs. Mean expression levels for AGO1-binding miRNAs were similar to that for all other miRNAs, at 7,279.36 counts vs 7,183.92 counts, respectively. In addition to *AGO1* mRNA, HEK293 cell line expressed two other mRNAs displaying non-specific sponge-like effect, *HSPA1B* with 77 interactions to 41 different microRNAs and *EEF1A1* with 50 interactions to 42 microRNAs. Similar to artificially over-expressed AGO1mRNA, *EEF1A1* also highly expressed in HEK293 cell line (>19K tpm), while another “sponge-like” mRNA *HSPA1B* had expression level less than 1 tpm.

A set of “sponge-like” mRNAs expressed in Huh7.5 cell line was entirely different. There were two protein-coding mRNAs, one for *AFP* - 47 interactions with 32 microRNAs and one for *APOB* - 47 interactions with 32 microRNAs, and two long-noncoding mRNAs, *MALAT1* with 47 interactions to 27 microRNAs and *XIST* with 55 interactions to 31 microRNAs. In coherence to expression levels of “sponge-like” mRNAs in HEK293 cell line, we observed different expression level for these mRNAs: *AFP* – more than 19K, *APOB* – 358 tpm, *XIST* – 202 tpm and *MALAT1* – 80 tpm, while the average expression level in Huh7.5 was – 69 tpm.

#### 3.2.2 Comparative analysis of microRNA expression levels and their mRNA interacting properties

To assess the role of microRNAs in the regulation of their target mRNAs, we studied two HEK293 and three Huh7.5 miRNA profiles retrieved from RNAseq datasets deposited in GEO (GSE75136 and GSE74014). For each cell line, only high-quality datasets with very high correlation of miRNA-specific expression levels were selected (Pearson’s correlation r >>0.99). For each miRNA, we analysed their cell-line specific levels of expression by R package “DeSeq2” in order to normalize miRNA expression, and compared these levels to the sets of experimentally detected microRNA-mRNA interactions retrieved from HEK293/CLASH and Huh7.5/ CLEAR-CLIP datasets MicroRNA was considered as expressed if it had expression levels of more than 3 counts (see Methods). Less than a quarter (23.5%) of 989 detected miRNAs was present in both cell lines (Figure 2E, Supplementary Table 2). Notably, many microRNAs expressed in the HEK293 (N = 205) and Huh7.5 (N = 194) cell lines then failed experimental detection as mRNA interacting molecules in CLASH or CLEAR-CLIP, respectively.

On the other hand, both CLASH and CLEAR-CLIP datasets included many mRNA-interacting microRNAs not detected in respective RNAseq datasets at all. On average, these microRNAs had relatively small amounts of interactions: 2.2+/-0.6 interacting partners for 197 microRNAs present in CLASH dataset, but absent in HEK293-based RNAseq, and 5.1+/-2.2 interacting partners for 168 miRNAs present in CLEAR-CLIP dataset but absent in Huh7.5-based RNAseq. For comparison, mean amounts of detected interactions across all microRNAs were at 55.8 +/-12.7 for 398 miRNAs of HEK293/CLASH and at 143.5 +/-28.5 for 542 miRNAs in Huh7.5/CLEAR-CLIP.

Next, for each of microRNAs we evaluated its cell-specific expression level and the amount of interactions in this cell line (Supplementary Figure 2). For each cell line, Spearman correlations levels were quite low, at 0.18 and 0.29 in HEK293 (N=335) and Huh7.5 (N=342), respectively. For each miRNA, we calculated the cell line-specific ratios (R) of its expression level to amount of detected interactions. The detailed analysis of this data allowed us to highlight two interesting types of miRNA. Type 1 comprised microRNAs with high expression level and relatively small amount of interactions with respective mRNAs. When the cut-offs for both R and expression levels were set as ranking at 90th percentile or higher, only 16 miRNAs for HEK293 (expression > 4418 and ratio > 252) and 12 miRNAs in Huh7.5 (expression > 6941 and ratio > 209) were classified as Type 1. Notably, eight Type 1 miRNAs were present in both cell lines examined.

Type 2 microRNAs were characterized by a low R and many detected interactions with mRNAs. When the cut-off for R was set as ranking at 10th percentile or lower, and amounts of interactions at 90^th^ percentile or higher, only 11 and 6 miRNAs for HEK293 (amount of interactions > 150 and ratio < 0.9) and Huh7.5 (amount of interactions > 165 and ratio < 2.5), were classified as Type 2, respectively. Unlike the Type 1 microRNAs, Type 2-specific sets from HEK293 and Huh7.5 did not overlap.

In order to evaluate whether these types of microRNAs are evolutionarily constrained, for all mature microRNAs from miRBase we calculated the mean of the phyloP conservative values in 20 vertebrates. The average cell line-specific phyloP scores for the Type 1 and Type 2 microRNAs were similar, at 0.99 and 0.95, respectively. Notably, these scores were higher than the average score value calculated for all known microRNAs (0.24) and the score values for all microRNAs that were identified as expressed or interacted in HEK293 or Huh7.5 cell lines (0.74 and 0.71, respectively). Notably, 80% of Top-100 miRbase microRNAs with the highest conservative phyloP scores were seen either as expressed or interacted (or both) in at least one of these two cell lines. On average, in HEK293 and Huh7.5 cells, these most conservative microRNAs had two times higher expression levels than less conservative expressed microRNAs (Supplementary Table 2). Overall, higher than average conservativeness of Type 1 and Type 2 microRNAs may point at the relative importance of their functions.

#### 3.2.3 Comparing cellular contexts for microRNA’s interactions

As expected, a majority of microRNAs were concordant in two cell lines: their expression levels and amounts of mRNA interactions were similar in both cellular contexts (Supplementary Figure 3A). Nevertheless, some miRNAs have demonstrated remarkable cell specificity in their ratios R (Supplementary Figure 3BC).

**Figure 3.**
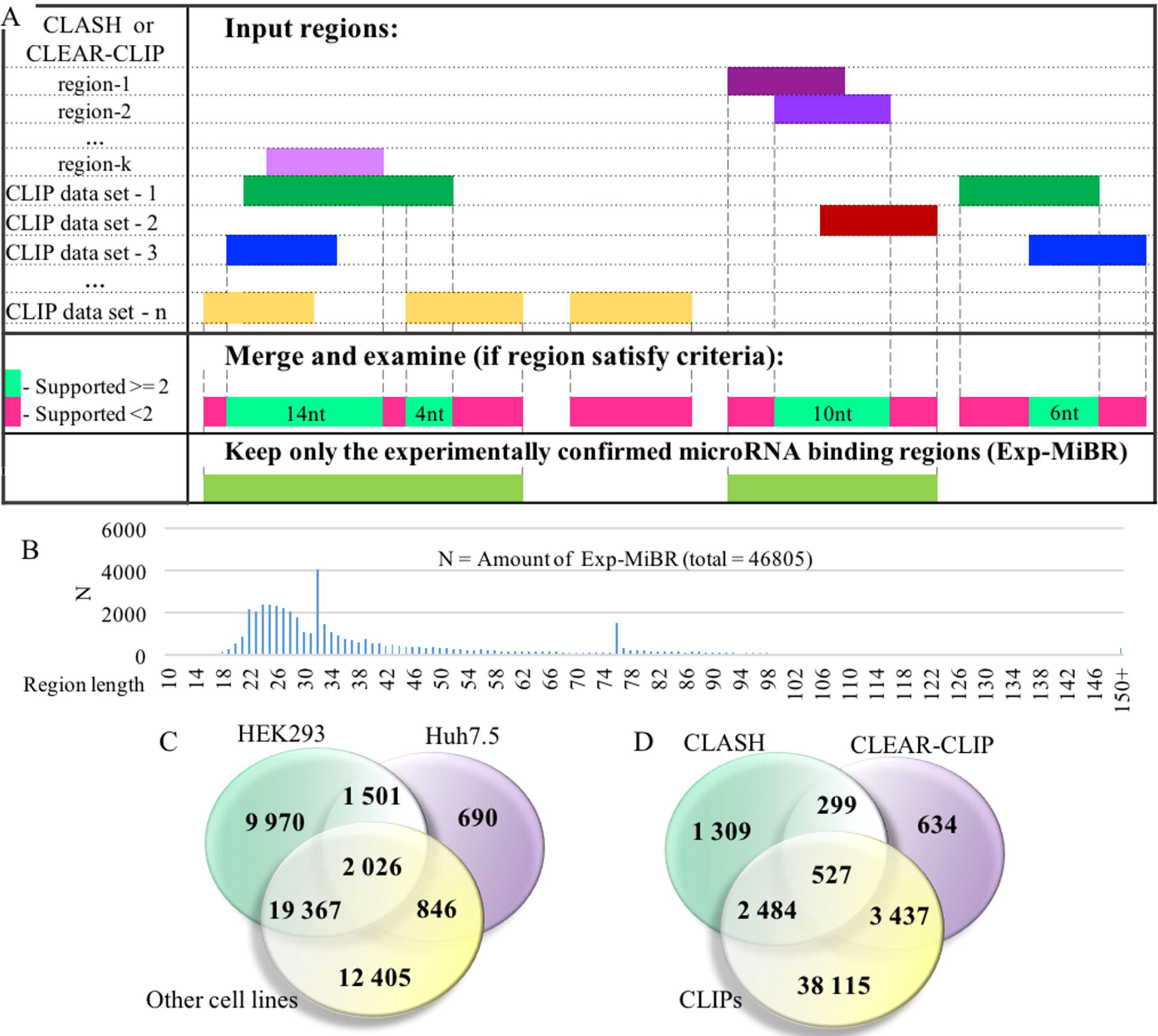
The detailed analysis of experimentally confirmed microRNA binding regions (Exp-MiBRs). **(A)** Validation of the Exp-MiBR by their independent occurrence in two or more datasets, or in two or more chimeric sequences from one dataset. **(B)** Exp-MiBRs: distribution of the lengths. On horizontal axis – the length of the Exp-MiBRs subsequence; on vertical axis – amounts of the detected Exp-MiBRs (N). **(C)** Venn diagram depicting tissue specificity of Exp-MiBRs detected in HEK293, Huh7.5 and all other cell lines **(D)**. Venn diagram depicting Exp-MiBRs detected in experiments employing three different types of identification techniques.

For 30 microRNAs, we detected high concordance between their expression level and amount of experimentally detected interactions. Eighteen of these miRNAs had higher expression and mRNA binding activity in Huh7.5 cell line, while for 12 remaining microRNA, both mRNA binding activity and expression level were higher in HEK293 cells (Supplementary Figure 3B). As an example, in Huh7.5 cell line, expression levels of MAPK1-repressing hsa-miR-194-5p (Kong et al., 2018) were 89 times higher than that in HEK293 cells; in Huh7.5 cells, this microRNA displayed 336 interactions, while in HEK293 it formed only 7 interactions. On the other hand, in HEK293, expression levels of lanosterol synthase suppressing hsa-miR-10a-5p (Kim et al., 2018) were 450 times higher than that in Huh7.5 cells; in HEK293 cells, this microRNA displayed 267 interactions, while in Huh7.5 it formed only 8 interactions. Such observations were expectable: microRNAs with higher expression level may be capable of the binding to a larger repertoire of targets.

Peculiarly, a total of microRNAs have performed in exactly opposite way: in cells with higher expression levels, these microRNAs displayed lesser amounts of interactions with their mRNA targets (Supplementary Figure 3C). For example, in Huh7.5 cell line, expression levels of hsa-miR-331-3p and hsa-miR-100-5p were at 1030 and 916 counts, respectively, while in HEK293 these miRNAs had 65 and 41 expression counts, respectively. However, in both cases, amounts of interactions in Huh7.5 cell line were lesser than that in HEK293 cell line, 47 versus 342 partners for hsa-miR-331-3p, and 1 versus 30 partners for hsa-miR-100-5p. To investigate if this phenomenon is due to the difference in the cell-specific expression levels of target genes, we performed an analysis of all these targets. This was, as well, not the case. As an example, only 21 out of 318 individual miRNA targets of hsa-miR-331-3p, were active in HEK293 cell line, but not detected in Huh7.5.

### 3.3 Analysis of expanded set of experimentally confirmed microRNA binding regions

Experimentally identified microRNA binding regions form a promising basis for further queries into the basics of the gene expression regulation, and lead to uncovering novel disease-causing mechanisms. To enhance a set of microRNA-mRNA interactions retrieved from CLASH and CLEAR-CLIP studies, we performed the database integration of the data collected in cross-linking with immunoprecipitation (CLIP) experiments that provide information about microRNA binding regions of target genes, but unable to identify mRNA-microRNAs pairings.

For this purpose, we collected data from 79 CLIP experiments, comprising 61 HITS-CLIP and 18 PAR-CLIP datasets covering 9 different cell lines, with a majority of these data obtained either in HEK293 (N=34 datasets) or Huh7.5 (N=19 datasets) cell lines (Supplementary Table 3). After combining CLIP datasets with the data of previously mentioned CLASH and CLEAR-CLIP studies, approximately 156,000 unique microRNA binding regions catalogued within mRNA targets.

At the next stage, the set of microRNA binding regions was cleaned up to include only these satisfying following criteria: (i) every position in this microRNA-binding subsequence is supported by evidence from at least two different datasets or two different chimeric sequences and (ii) the length of at least 10nt (Figure 3A, Supplementary Table 4). MiRNA-binding subsequences of this kind (N = 46,805) formed a dataset of experimentally confirmed microRNA binding regions (Exp-MiBR). In this dataset, each Exp-MiBR record includes following attributes: genomic coordinates, gene name, type of mRNA part, list of GEO GSM IDs for experiments which support this microRNA interaction, cellular context, and the list of interacting microRNAs (if accessible). The criteria for inclusion of individual microRNA-binding regions in Exp-MiBR database are justified by analysis presented in Supplementary Table 5.

Exp-MiBR subsequences (N = 46,805) were mapped to approximately 15,000 human genes. About one-half of Exp-MiBRs (48%) were located in 3’UTRs, 24% in a coding part, 10% in introns and 6% in intergenic parts. Remaining 10% of the Exp-MiBRs were mapped to non-coding RNAs, being matched to either exonic or intronic regions of these loci.

Approximately 68% of Exp-MiBRs were 20-40 nt in size, closely matching the mean length (33 nt) for all input miRNA-binding regions from CLIPs, CLASH and CLEAR-CLIP data (Figure 3B). The second peak in size distribution of Exp-MiBRs was at 75 to 80 nt, being predominantly comprised (86%) of miRNA-interacting region extracted from CLEAR-CLIP dataset. While the sizes of 99% of the Exp-MiBRs were smaller than 150nt, a few Exp-MiBRs were much longer than that, while remaining supported by many experiments. The longest Exp-MiBR of 631 nt was formed by the regions confirmed as microRNA-interacting in 54 different experiments in nine different cell lines. In addition, there were a few Exp-MiBRs located closely to each other. Such clusters of Exp-MiBRs with many interacting microRNAs do not display a tendency to any particular region of mRNA, as they may be present in CDS, 3’UTR, 5’UTR or intergenic regions. As an example, chromosome 2 contains a cluster of Exp-MiBRs covering an area of approximately 1.5 kb in size, which is located between the loci of RNA5-8SP5 and MIR663B genes. According to CLASH and CLEAR-CLIP studies, this cluster of Exp-MiBRs interacts with 52 different miRNAs (Supplementary Figure 4, Supplementary Table 6).

### 3.4 Tissue-specific and housekeeping microRNA binding regions

To characterize Exp-MiBRs further, we analyzed their tissue specificity. Most CLIP experiments were performed either in HEK293 (43%) or in Huh7.5 (24%) cells, while the rest of the CLIP data were collected in HeLa, HFF, BC-1, BC-3, EF3D, LCL35 or LCL cells. In HEK293 cells, we found approximately 9,900 unique MiBRs, while analysis of Huh7.5 cells yielded 690 tissue-specific interacting regions (Figure 3C). Larger amounts of Exp-MiBRs in HEK293 as compared to that Huh7.5 cells may be explained either by better coverage of HEK293 transcriptome by various CLIPs (Supplementary Table3), or by intrinsic cell-specific features of miRNA interactomes.

Interestingly, some Exp-MiBRs were observed a majority of studied cells, possibly reflecting a housekeeping function of these interactions. Approximately 1% of all Exp-MiBRs were found in seven or more cell lines. The functional roles of 351 ubiquitous Exp-MiBRs were investigated using Panther software. The GO analysis showed enrichment of genes participating in cellular process of cell cycle (FC 3.17; p-value 1e10-8) and in molecular function of nucleic acid binding (FC 1.75; p-value 5e10-4).

### 3.5 Mitochondrial regulation by microRNA

An analysis of Exp-MiBRs revealed that these microRNA interacting sequences cover 86% of the mitochondrial genome, including 35 out of 37 mitochondrial genes. Mitochondrial Exp-MiBRs (N = 37) were found in all nine investigated cell lines, with each Exp-MiBR discovered, on average, in 11 independent experiments. In total, we identified 182 miRNAs that bound various mitochondrial RNAs, with two mitochondrial regions binding 107 out of 182 miRNAs.

## 4 Discussion

Experimental identification of microRNA binding regions is an important prerequisite for querying into the basics of the gene expression regulation, and for uncovering novel disease-causing mechanisms. To date, only two sequencing-based experimental datasets describing full miRNA-mRNA interactomes of human cells, CLASH and CLEAR-CLIP, are available. In both studies, the primary goal was to develop and optimize the experimental protocol itself, while identifying miRNA-mRNA interactions in a particular cell line grown under different conditions. Although these techniques provide unique window into miRNA targeting, they are not free of limitations, which precludes determining of entire miRNA-mRNA interactome. Nevertheless, intersecting CLASH and CLEAR-CLIP datasets allowed us to detect much larger set of validated interactions than may be expected of two randomly-generated datasets.

Typically, miRNA-mRNA interaction networks built in silico with an aid of one or another miRNA prediction tool include thousands of mRNA targets. In our study, we attempted to paint a holistic picture of human miRNA-mRNA interactome by comparing the entries from experimentally collected datasets describing miRNA binding activity to the data describing expression data. Interestingly, we found that more than half of mRNA transcripts do not bind to any miRNAs present in the same cellular environment, while 1-2% of human transcripts interact with nine or more miRNAs, thus, displaying a similar to sponge-like activity (Thomson and Dinger, 2016). Remarkably, miRNA-mRNA sponge-like interactions were cell-lines specific, with very little overlap identified. In HEK293 cells, the most prominent sponge-like activity resultant in 77 different miRNA interactions was detected for *AGO1* mRNA, which had been initially overexpressed according to the CLASH protocol. Two other “sponge-like” mRNAs *HSPA1B* and *EEF1A1* in HEK293 cell line formed 77 and 50 interactions respectively.

This amount of interactions is comparable to that of a well-known circular RNA with sponge properties, Cdr1as (74 predicted sites) (Xu et al., 2015). In Huh7.5 cells, the set of RNAs with “sponge-like” activities included many noncoding RNAs, including *MALAT1* and *XIST*. It is peculiar that some Huh7.5–specific sponge-like RNAs, including these for alpha-fetoprotein (*AFP*) (Parpart et al., 2014) and *APOB* (Bi et al., 2014) were previously described as biomarkers of liver carcinoma, a tissue of origin for Huh7.5 cell line.

Some miRNAs expressed at relatively high levels were not among RNA interactors at all. About a hundred of such non-interacting miRNAs were present in both studied cell lines. There is a possibility that the natural targets for these microRNAs are either not expressed in studied cellular contexts, or that they have no targets at all. In total, only 232 microRNAs had at least one interaction in each of studied cell lines.

For individual miRNAs, levels of their expression have no bearing on amounts of interactions they display, possibly reflecting difference in their functions depending on the cellular context. As an example, we revealed that, in Huh7.5 cell line, miR-423-3p is abundant but displays only a few interactions, while in HEK293 cell line the same miRNA forms more than two hundred interactions and expressed at the quite low level. These observations complement previous findings of Mullokandov and colleagues (Mullokandov et al., 2012), who have shown that the binding activity of some highly expressed miRNAs may be weakened by either high target-to-miRNA ratio or the relocation of this miRNA to the nucleus. Future studies are required for to investigate how RNA binding properties of individual miRNAs may change in response to regulation by context-dependent extrinsic or intrinsic factors.

Augmenting CLASH and CLEAR-CLIP datasets with additional 79 CLIP datasets provided us with information about microRNA footprints resulted in many thousands of experimentally confirmed microRNA binding regions (Exp-MiBR) present in both coding and noncoding regions of RNA loci. At least some Exp-MiBR are tissue-specifics, in agreement with Clark and colleagues, who revealed the differences in the microRNA targetomes across tissues (Clark et al., 2014).

In addition to chromosomes, many Exp-MiBRs map to mitochondrial DNA, where they are quite abundant. Previous studies showed four mitochondrial regions with high degree of homology to microRNAs, namely, hsa-miR-4461 (chrM: 10690–10712), hsa-miR-4463 (chrM: 13050– 13068), hsa-miR-4484 (chrM: 5749–5766) and hsa-miR-4485 (chrM: 2562–2582) (Sripada et al., 2012). Two of these regions, that encode mitochondrial *ND4L* and 16S rRNA genes, were also highly interacting Exp-MiBRs, with 70 and 63 cognate miRNAs, respectively, all confirmed in nine different cell lines. In both cases, previously identified cognate miRNAs hsa-miR-4461 and hsa-miR-4485 were among confirmed interactors. Our study expands the coverage of mitochondrial genome by various miRNA-interacting regions to 86% of its lengths. Altogether, these findings support the notion that miRNA– mRNA interactions take place in a variety of cellular compartments, including mitochondria (Ni and Leng, 2015).

Analysis of the landscape of microRNA-mRNA human interactions, which we derived from both direct microRNA-mRNA interactions experimentally defined in HEK293 and Huh7.5 cell lines, along with microRNA and mRNA expression data highlight complexity of human microRNA-mRNA interactome. For individual miRNAs, levels of their expression have no bearing on amounts of interactions they display, possibly reflecting difference in their functions depending on the cellular context. In this article, we found that while only 1-2% of human genes were the most regulated by microRNAs, a few cell line specific RNAs display a similar to sponge effect: *EEF1A1* and *HSPA1B* in HEK293 and *AFP, APOB* and *MALAT1* genes in Huh7.5 cell lines. Some miRNAs might be expressed at relatively low levels, and interact with many mRNAs. On the other hand, there is a set of microRNAs expressed at a very high level and interacting with only a few mRNAs, thus, indeed, regulating expression of their targets in a specific manner. Notably, microRNAs are capable of switching between these two modes of action, depending on cellular context. The question of the biological significance of these two miRNA groups is still open. CLASH and/or CLEAR-CLIP coverage of additional cell lines is warranted. It is notable, however, that the presence of miRNA groups, one with a low expression level and a high number of interactions, and one with opposite characteristics, was detected in both cell lines profiled.

We have also established a collection of reliable microRNA binding regions that we systematically extracted in course of analysis of 79 CLIP datasets, which is available at http://score.generesearch.ru/services/mirna/. The promise of microRNAs as potential means for diagnostics and therapy got expanded with a number of loss-of-function and, recently, the case of disease-causing gain-of-function mutation in particular microRNA (Grigelioniene et al., 2019). We believe that the results of our efforts in mapping the human miRNA-mRNA interactome may be useful in untangling molecular underpinnings of hereditary and acquired diseases that involve interactions.

## 6 Author Contributions

MS and OP designed the study and carried out the research. AB contributed to the discussion of the results. OP and AB wrote the paper. All authors read and approved the final manuscript.

## List of abbreviations

AGO: Argonaute
CDS: Coding DNA sequence
CLASH: Crosslinking, ligation and sequencing of hybrids technique
CLEAR-CLIP: covalent ligation of endogenous Argonaute-bound RNA-CLIP technique
CLIP-UV: crosslinking and immunoprecipitation technique
Exp-MiBRs: experimentally confirmed microRNA binding regions
HITS-CLIP: High-throughput sequencing of RNA isolated by crosslinking immunoprecipitation
iCLIP: individual-nucleotide resolution Cross-Linking and ImmunoPrecipitation
PAR-CLIP: Photoactivatable-Ribonucleoside-Enhanced Immunoprecipitation
UTR: Untranslated region

## 9 Acknowledgments

We thank Andrey Marakhonov and members of the Skoblov laboratory for helpful discussions.

## References

Agarwal, V., Bell, G. W., Nam, J. W., Bartel, D. P. (2015). Predicting effective microRNA target sites in mammalian mRNAs. Elife. 4 doi: 10.7554/eLife.05005.

Artcibasova, A. V., Korzinkin, M. B., Sorokin, M. I., Shegay, P. V., Zhavoronkov, A. A., Gaifullin, N., et al. (2016). MiRImpact, a new bioinformatic method using complete microRNA expression profiles to assess their overall influence on the activity of intracellular molecular pathways. Cell Cycle. 15(5), 689–698. doi: 10.1080/15384101.2016.1147633.

Bandiera, S., Pernot, S., El Saghire, H., Durand, S. C., Thumann, C., Crouchet, E., et al. (2016). Hepatitis C Virus-Induced Upregulation of MicroRNA miR-146a-5p in Hepatocytes Promotes Viral Infection and Deregulates Metabolic Pathways Associated with Liver Disease Pathogenesis. J. Virol. 90(14), 6387–6400. doi: 10.1128/JVI.00619-16.

Bartel, DP. (2004). MicroRNAs: genomics, biogenesis, mechanism, and function. Cell. 116(2), 281–97.

Bi, Y., He, Y., Huang, J., Su, Y., Zhu, G. H., Wang, Y., et al. (2014). Functional characteristics of reversibly immortalized hepatic progenitor cells derived from mouse embryonic liver. Cell. Physiol. Biochem. 34(4), 1318–38. doi: 10.1159/000366340.

Bray, N. L., Pimentel, H., Melsted, P., Pachter, L. (2016). Near-optimal probabilistic RNA-seq quantification. Nat. Biotechnol. 34(5), 525–7. doi: 10.1038/nbt.3519.

Clark, P. M., Loher, P., Quann, K., Brody, J., Londin, E. R., Rigoutsos, I. (2014). Argonaute CLIP-Seq reveals miRNA targetome diversity across tissue types. Sci Rep. 4, 5947. doi: 10.1038/srep05947.

Corcoran, D. L., Georgiev, S., Mukherjee, N., Gottwein, E., Skalsky, R. L., Keene, J. D., et al. (2011). PARalyzer: definition of RNA binding sites from PAR-CLIP short-read sequence data. Genome Biol. 12(8), R79. doi: 10.1186/gb-2011-12-8-r79.

Friedman, R. C., Farh, K. K. H., Burge, C. B., Bartel, D. P. (2009). Most mammalian mRNAs are conserved targets of microRNAs. Genome Res. 19(1), 92–105. doi: 10.1101/gr.082701.108.

Grigelioniene, G., Suzuki, H. I., Taylan, F., Mirzamohammadi, F., Borochowitz, Z. U., Ayturk, U. M., et al. (2019). Gain-of-function mutation of microRNA-140 in human skeletal dysplasia. Nature medicine. 1. doi: 10.1038/s41591-019-0353-2.

Grimson, A., Farh, K. K. H., Johnston, W. K., Garrett-Engele, P., Lim, L. P., Bartel, D. P. (2007). MicroRNA targeting specificity in mammals: determinants beyond seed pairing. Mol. Cell. 27(1), 91–105. doi: 10.1016/j.molcel.2007.06.017.

Gumienny, R, and Zavolan, M. (2015). Accurate transcriptome-wide prediction of microRNA targets and small interfering RNA off-targets with MIRZA-G. Nucleic Acids Res. 43(3), 1380–91. doi: 10.1093/nar/gkv050.

He, L, and Hannon, GJ. (2004). MicroRNAs: small RNAs with a big role in gene regulation. Nat. Rev. Genet. 5(7), 522–31. doi: 10.1038/nrg1379.

Heberle, H., Meirelles, G. V., da Silva, F. R., Telles, G. P., Minghim, R. (2015). InteractiVenn: a web-based tool for the analysis of sets through Venn diagrams. BMC Bioinformatics. 16, 169. doi: 10.1186/s12859-015-0611-3.

Helwak, A., Kudla, G., Dudnakova, T., Tollervey, D. (2013). Mapping the human miRNA interactome by CLASH reveals frequent noncanonical binding. Cell. 153(3), 654–65. doi: 10.1016/j.cell.2013.03.043.

Hu, B., Yang, Y. C. T., Huang, Y., Zhu, Y., Lu, Z. J. (2016). POSTAR: a platform for exploring post-transcriptional regulation coordinated by RNA-binding proteins. Nucleic Acids Res. 45(D1), D104–D114. doi: 10.1093/nar/gkw888.

Jonas, S, and Izaurralde, E. (2015). Towards a molecular understanding of microRNA-mediated gene silencing. Nat. Rev. Genet. 16(7), 421–33. doi: 10.1038/nrg3965.

Karolchik, D., Hinrichs, A. S., Furey, T. S., Roskin, K. M., Sugnet, C. W., Haussler, D., et al. (2004). The UCSC Table Browser data retrieval tool. Nucleic Acids Res. 32(Database issue), D493–6. doi: 10.1093/nar/gkh103.

Kim, J. E., Hong, J. W., Lee, H. S., Kim, W., Lim, J., Cho, Y. S., et al. (2018). Hsa-miR-10a-5p downregulation in mutant UQCRB-expressing cells promotes the cholesterol biosynthesis pathway. Sci Rep. 8(1), 12407. doi: 10.1038/s41598-018-30530-6.

Kong, Q., Zhang, S., Liang, C., Zhang, Y., Kong, Q., Chen, S., et al. (2018). LncRNA XIST functions as a molecular sponge of miR-194-5p to regulate MAPK1 expression in hepatocellular carcinoma cell. J. Cell. Biochem. 119(6), 4458–4468. doi: 10.1002/jcb.26540.

Kozomara, A, and Griffiths-Jones, S. (2014). miRBase: annotating high confidence microRNAs using deep sequencing data. Nucleic Acids Res. 42(Database issue), D68–73. doi: 10.1093/nar/gkt1181.

Li, Y., and Zhang, Z. (2014). Potential microRNA-mediated oncogenic intercellular communication revealed by pan-cancer analysis. Scientific reports. 4, 7097. doi: 10.1038/srep07097.

Li, Y., Liang, C., Wong, K. C., Jin, K., Zhang, Z. (2014). Inferring probabilistic miRNA–mRNA interaction signatures in cancers: a role-switch approach. Nucleic acids research. 42(9), e76–e76. doi: 10.1093/nar/gku182.

Licatalosi, D. D., Mele, A., Fak, J. J., Ule, J., Kayikci, M., Chi, S. W., et al. (2008). HITS-CLIP yields genome-wide insights into brain alternative RNA processing. Nature. 456(7221), 464–9. doi: 10.1038/nature07488.

Lu, Y, and Leslie, CS. (2016). Learning to Predict miRNA-mRNA Interactions from AGO CLIP Sequencing and CLASH Data. PLoS Comput. Biol. 12(7), e1005026. doi: 10.1371/journal.pcbi.1005026.

Luna, J. M., Scheel, T. K., Danino, T., Shaw, K. S., Mele, A., Fak, J. J., et al. (2015). Hepatitis C virus RNA functionally sequesters miR-122. Cell. 160(6), 1099–110. doi: 10.1016/j.cell.2015.02.025.

Moore, M. J., Scheel, T. K., Luna, J. M., Park, C. Y., Fak, J. J., Nishiuchi, E., et al. (2015). miRNA-target chimeras reveal miRNA 3’-end pairing as a major determinant of Argonaute target specificity. Nat Commun. 6, 8864. doi: 10.1038/ncomms9864.

Moore, M. J., Zhang, C., Gantman, E. C., Mele, A., Darnell, J. C., Darnell, R. B. (2014). Mapping Argonaute and conventional RNA-binding protein interactions with RNA at single-nucleotide resolution using HITS-CLIP and CIMS analysis. Nat Protoc. 9(2), 263–93. doi: 10.1038/nprot.2014.012.

Mullokandov, G., Baccarini, A., Ruzo, A., Jayaprakash, A. D., Tung, N., Israelow, B., et al. (2012). High-throughput assessment of microRNA activity and function using microRNA sensor and decoy libraries. Nat. Methods. 9(8), 840–6. doi: 10.1038/nmeth.2078.

Murakawa, Y., Hinz, M., Mothes, J., Schuetz, A., Uhl, M., Wyler, E., et al. (2015). RC3H1 post-transcriptionally regulates A20 mRNA and modulates the activity of the IKK/NF-κB pathway. Nat Commun. 6, 7367. doi: 10.1038/ncomms8367.

Ni, WJ, and Leng, XM. (2015). Dynamic miRNA-mRNA paradigms: New faces of miRNAs. Biochem Biophys Rep. 4, 337–341. doi: 10.1016/j.bbrep.2015.10.011.

Parpart, S., Roessler, S., Dong, F., Rao, V., Takai, A., Ji, J., et al. (2014). Modulation of miR-29 expression by α-fetoprotein is linked to the hepatocellular carcinoma epigenome. Hepatology. 60(3), 872–83. doi: 10.1002/hep.27200.

Plotnikova, OM, and Skoblov, MY. (2018). Efficiency of the miRNA-mRNA Interaction Prediction Programs. Mol. Biol. (Mosk.). 52(3), 543–554. doi: 10.7868/S0026898418030187.

Riffo-Campos, Á., Riquelme, I., Brebi-Mieville, P. (2016). Tools for Sequence-Based miRNA Target Prediction: What to Choose?. Int J Mol Sci. 17(12) doi: 10.3390/ijms17121987.

Sætrom, P., Heale, B. S., Snøve Jr, O., Aagaard, L., Alluin, J., Rossi, J. J. (2007). Distance constraints between microRNA target sites dictate efficacy and cooperativity. Nucleic Acids Res. 35(7), 2333–42. doi: 10.1093/nar/gkm133.

Selbach, M., Schwanhäusser, B., Thierfelder, N., Fang, Z., Khanin, R., Rajewsky, N. (2008). Widespread changes in protein synthesis induced by microRNAs. Nature. 455(7209)б 58–63. doi: 10.1038/nature07228.

Sripada, L., Tomar, D., Prajapati, P., Singh, R., Singh, A. K., Singh, R. (2012). Systematic analysis of small RNAs associated with human mitochondria by deep sequencing: detailed analysis of mitochondrial associated miRNA. PLoS ONE. 7(9), e44873. doi: 10.1371/journal.pone.0044873.

Steinkraus, B. R., Toegel, M., Fulga, T. A. (2016). Tiny giants of gene regulation: experimental strategies for microRNA functional studies. Wiley Interdiscip Rev Dev Biol. 5(3), 311–62. doi: 10.1002/wdev.223.

Thomson, DW, and Dinger, ME. (2016). Endogenous microRNA sponges: evidence and controversy. Nat. Rev. Genet. 17(5), 272–83. doi: 10.1038/nrg.2016.20.

Uhlmann, S., Mannsperger, H., Zhang, J. D., Horvat, E. Á., Schmidt, C., Küblbeck, M., et al. (2012). Global microRNA level regulation of EGFR-driven cell-cycle protein network in breast cancer. Mol. Syst. Biol. 8б 570. doi: 10.1038/msb.2011.100.

Wang, K., Li, M., Hakonarson, H. (2010). ANNOVAR: functional annotation of genetic variants from high-throughput sequencing data. Nucleic Acids Res. 38(16), e164. doi: 10.1093/nar/gkq603.

Weiss, CN, and Ito, K. (2017). A Macro View of MicroRNAs: The Discovery of MicroRNAs and Their Role in Hematopoiesis and Hematologic Disease. Int Rev Cell Mol Biol. 334, 99–175. doi: 10.1016/bs.ircmb.2017.03.007.

Wissink, E. M., Fogarty, E. A., Grimson, A. (2016). High-throughput discovery of post-transcriptional cis-regulatory elements. BMC Genomics. 17, 177. doi: 10.1186/s12864-016-2479-7.

Xu, H., Guo, S., Li, W., Yu, P. (2015). The circular RNA Cdr1as, via miR-7 and its targets, regulates insulin transcription and secretion in islet cells. Sci Rep. 5, 12453. doi: 10.1038/srep12453.

Yang, H, and Wang, K. (2015). Genomic variant annotation and prioritization with ANNOVAR and wANNOVAR. Nat Protoc. 10(10), 1556–66. doi: 10.1038/nprot.2015.105.

